# HIF-dependent Neuropeptide Y Receptor Y1 and Y5 expression sensitizes hypoxic cells to NPY stimulation

**DOI:** 10.1101/2021.07.14.452372

**Authors:** Philip J. Medeiros, Sydney Pascetta, Sarah Kirsh, Baraa K. Al-Khazraji, James Uniacke

**Author notes:** Current address: Department of Kinesiology and Physical Education, Wilfrid Laurier University, Waterloo, Ontario, Canada (Philip J. Medeiros). Corresponding author: Department of Molecular and Cellular Biology, University of Guelph, 50 Stone Road East, Guelph, Ontario, Canada N1G 2W1.

## Abstract

Neuropeptide Y (NPY) is an abundant neurohormone in the central and peripheral nervous system involved in feeding behavior, energy balance, nociception, and anxiety. Several NPY receptor (NPYR) subtypes display elevated expression in many cancers including in breast cancer where this is exploited for imaging and diagnosis. Here, we show that NPY1R and NPY5R mRNA abundance is induced by hypoxia in a Hypoxia Inducible Factor (HIF)-dependent manner in breast cancer cell lines MCF7 and MDA-MB-231. The HIFs bind to several genomic regions upstream of the *NPY1R* and *NPY5R* transcription start sites. The MAPK/ERK pathway is activated more rapidly upon NPY5R stimulation in hypoxic cells compared to normoxic cells. This pathway requires IGF1R activity in normoxia, but not in hypoxic cells where they display resistance to the radiosensitizer and IGF1R inhibitor AG1024. Hypoxic cells proliferate and migrate more when stimulated with NPY relative to normoxic cells, with a more robust response observed with a Y5-specific agonist. Our data suggest that hypoxia induced NPYRs render hypoxic cells more sensitive to NPY stimulation. Considering that breast tissue receives a constant supply of NPY, and hypoxia is a common feature of the tumor microenvironment, breast tumors are the perfect storm for hyperactive NPYR. This study not only highlights a new relationship between the HIFs and NPYR expression and activity, but may inform the use of chemotherapeutics targeting NPYRs and hypoxic cells.

## INTRODUCTION

The 36-amino acid Neuropeptide Y (NPY) neurohormone family is broadly distributed throughout the central and peripheral nervous systems and is the most abundant neuropeptide in the brain (1,2). Within the central nervous system, NPY is involved in the regulation of feeding behavior, energy balance, nociception, and anxiety. In the peripheral nervous system, the pleiotropic actions of NPY include regulation of vascular tone, chemotaxis of endothelial and vascular smooth muscle cells, leukocyte migration, and angiogenesis (1,3). Within the periphery, NPY is primarily stored and released from sympathetic nerves of the autonomic branch of the peripheral nervous system and is released following high frequency neuronal stimulation (3).

There are five well-defined NPY receptor (NPYR) subtypes in mammals: Y1, Y2, Y4, Y5 and Y6. However, NPY6R is encoded by a non-functional pseudogene in primates, and NPY4R exhibits low affinity for NPY (4). Thus, NPY1R, NPY2R and NPY5R are the most biologically significant subtypes in humans (3,5). NPYRs are class I rhodopsin-like family of G-protein coupled receptors (GPCRs) and mediate their effects through a trimeric inhibitory GTP-binding protein (Gi/o) (3). All NPYRs decrease adenylyl cyclase activity, but additional receptor-mediated signaling pathways are generally less understood based on subtype- and tissue-specific interactions (3,6). Sharing 60% of the same amino acids (7), NPY1R and NPY5R have been implicated in activating the MAPK pathway in cardiomyocytes (8), breast cancer cells (9,10), and in HEK293 cells through transactivation with Insulin-like Growth Factor-1 Receptor (IGF1R) (11).

A growing body of evidence has linked NPYR expression with neoplastic transformation. NPY1R and NPY5R expression has been reported in several types of endocrine and epithelial malignancies, such as ovarian, prostate, breast, and neural crest tumors relative to normal tissue (10,12–16). Approximately 85% of primary human breast carcinomas exhibit elevated NPY1R expression (10). Further, many commonly used breast cancer cell lines such as MCF7 and MDA-MB-231 express NPY1R and NPY5R (10). Increased NPY5R expression in these cells leads to clustering of NPYR in the plasma membrane. Breast tissue is highly innervated by the sympathetic nervous system, thus it is provided with a large supply of NPY ligand. High local NPY concentration in combination with increased NPYR clustering leads to heightened NPYR activity (9,10). NPY1R and NPY5R stimulation promotes cellular proliferation, migration, and angiogenesis in breast cancer models (9,17,18). These observations not only highlight these NPYRs as markers of cancer, but potential targets of therapeutic intervention. Since epithelial malignancies such as breast carcinomas form into solid tumors, it is important to consider how the tumor microenvironment affects the progression of these cancers and the regulation of the NPYRs.

Hypoxia is a hallmark of solid tumors where oxygen consumption exceeds supply due to aberrant blood vessel formation (19). Tumor hypoxia leads to resistance to chemo- and radiation therapy, and promotes metastasis (20). The hypoxia inducible factors (HIFs) are central regulators of the transcriptional response to hypoxia. These heterodimeric transcription factors are linked to cancer progression and consist of an oxygen-regulated α-subunit (HIF-1α or HIF-2α) and a constitutively expressed β-subunit (HIF-1β) (21,22). The hypoxic stabilization of both HIFs leads to angiogenesis, metabolic reprogramming, immortalization, evasion of apoptosis, migration and invasion, generation of cancer stem cells, and chemo- and radiotherapy resistance. HIF-1α and HIF-2α share most of their transcription targets but do have distinct set of gene targets (23). Cancer cells can exploit HIF-mediated changes during neoplastic transformation to accelerate carcinogenesis, leading to a more aggressive tumor phenotype (19). In Ewing sarcoma cells, hypoxia shifts NPY activity from NPY1R/Y5R-mediated cell death in physiological conditions to NPY2R/Y5R-driven cell proliferation, migration, and angiogenesis (24).

Here, we investigate the relationship between NPYR signaling, hypoxia, and the cell migration and proliferation of breast cancer cell lines MCF7 and MDA-MB-231. Our data show that NPY1R and NPY5R mRNA is induced in hypoxia in a HIF-dependent manner. Further, the HIFs bind to several regions upstream of the *NPY1R* and/or *NPY5R* transcription start site (TSS). We show that NPY stimulation activates the MAPK/ERK pathway through IGF1R, but that this pathway is resistant to IGF1R inhibition in hypoxia. The induction of NPY1R and NPY5R sensitizes hypoxic cells to NPY ligand by activating MAPK/ERK more rapidly, and producing more cell migration and proliferation, relative to normoxic cells. Our data highlight that hypoxic cancer cells in an environment with high local NPY, such as the breast, may be resistant to radiosensitizers that inhibit IGF1R, and are strong candidates for NPYR antagonist therapy.

## RESULTS

### NPY1R and NPY5R are induced by hypoxia at the mRNA and protein level

MCF7 and MDA-MB-231 cells were incubated in hypoxia (1% O_2_) for 24 h and RNA was collected at 0 h, 1 h, 3 h, 6 h, 12 h, and 24 h. The temporal induction of NPY1R and NPY5R mRNA was measured via qRT-PCR. We selected this time frame because the major hypoxic gene expression pathway, HIF-mediated transcription, is only fully active at 24 h with each HIF-α isoform peaking at different times. HIF-1α protein peaks within 2 h of hypoxic exposure, while HIF-2α requires 24 h to achieve peak levels (Holmquist-Mengelbier L, et al. (2006)). In MCF7, the NPY1R displayed a significant 2.03 ± 0.09-fold increase in mRNA abundance after 1 h of hypoxia, and a significant 3.92 ± 0.2-fold increase after 3 h before returning to normoxic levels (time 0) after 24 h (**Fig. 1A**). In MDA-MB-231, NPY1R mRNA abundance peaked at a significant 2.9 ± 0.16-fold increase after 24 h of hypoxia relative to 0 h (**Fig. 1B**). The NPY5R mRNA abundance peaked at a significant 4.74 ± 0.13-fold increase at 3 h in MCF7 cells (**Fig. 1C**) and 3.93 ± 0.29 at 1 h in MDA-MB-231 cells (**Fig. 1D**) relative to 0 h. While the NPY1R and NPY5R mRNA abundance returned to baseline levels by 24 h in MCF7 cells (**Fig. 1A and C**), it remained at 2-3-fold baseline levels after 24 h in MDA-MB-231 (**Fig. 1B and D**).

**Figure 1.**
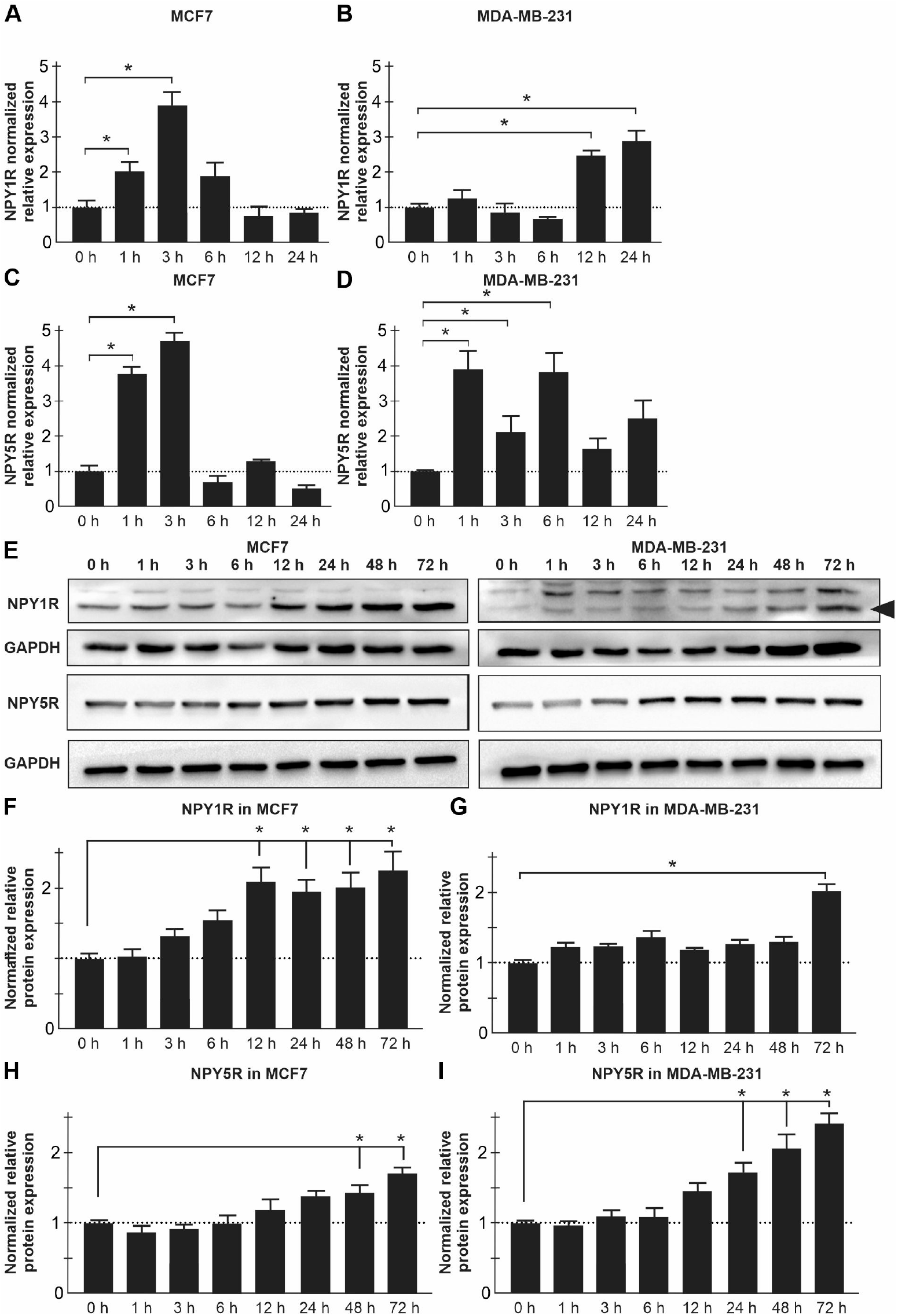
NPY1R and NPY5R are induced by hypoxia at the mRNA and protein level. (A-B) NPY1R mRNA and (C-D) NPY5R mRNA levels were measured by qRT-PCR in MCF7 (A and C) and MDA-MB-231 (B and D) breast cancer cells in a hypoxic (1% O_2_) time course of 0 h, 1 h, 3 h, 6 h, 12 h, and 24 h. Data normalized to endogenous control genes *RPLP0* and *RPL13A* and made relative to time 0 h (dotted line). (E) NPY1R (black arrow) and NPY5R protein levels measured in MCF7 and MDA-MB-231 cells in a hypoxic time course of 0 h, 1 h, 3 h, 6 h, 12 h, 24 h, 48 h, and 72 h. GAPDH was used as a loading control. Quantification by densitometry of western blots in (E) using ImageJ software for NPY1R in MCF7 (F) and MDA-MB-231 (G) cells, and NPY5R in MCF7 (H) and MDA-MB-231 (I) cells. Data (n = 3), mean ± s.e.m. * represents p < 0.05 using one-way ANOVA and Tukey’s HSD post-hoc test.

We next verified whether NPY1R and NPY5R hypoxic protein expression was consistent with the observations at the mRNA level. In both cell lines, NPY1R and NPY5R protein levels displayed significant increases with peaks that were delayed by several hours from the mRNA abundance data in Fig. 1. The NPY1R protein levels significantly increased by 2.11 ± 0.11-fold at 12 h of hypoxic exposure in MCF7 cells (**Fig. E-F**), and by 2.04 ± 0.05-fold at 72 h in MDA-MB-231 relative to 0 h (**Fig. 1E and G**). The NPY5R protein levels displayed significant increases in the 48-72 h range in MCF7 cells with a peak increase of 1.72 ± 0.05-fold at 72 h of hypoxic exposure relative to 0 h (**Fig. 1E and H**). In MDA-MB-231, NPY5R protein levels displayed significant increases in the 24-72 h range with a peak of 2.44 ± 0.08-fold at 72 h of hypoxic exposure relative to 0 h (**Fig. 1E and I**). These data suggest that NPY1R and NPY5R expression is induced in a hypoxia-dependent manner in MCF7 and MDA-MB-231 cells.

### A hypoxia mimetic that stabilizes the HIFs increases the mRNA abundance of NPY1R and NPY5R

To test whether an increase in HIF stability was responsible for the observed hypoxic increase in NPY1R and NPY5R mRNA abundance, we treated cells with Dimethyloxalylglycine (DMOG). DMOG is a cell permeable prolyl-4-hydroxylase inhibitor, which mimics hypoxia by preventing the hydroxylation of the HIF-α subunits leading to their stabilization. MCF7 and MDA-MB-231 cells were treated with 1 mM DMOG for 30 min, 1 h and 2 h followed by RNA collection and qRT-PCR. In MCF7, DMOG treatment produced a peak increase of 2.0 ± 0.06-fold for NPY1R mRNA at 1 h (**Fig. 2A**), and 2.41 ± 0.09-fold for NPY5R mRNA at 2 h relative to 0 h (**Fig. 2B**). In MDA-MB-231, NPY1R and NPY5R mRNA levels both peaked at 2 h of DMOG treatment with increases of 1.88 ± 0.12-fold and 6.95 ± 0.44-fold, respectively, relative to 0 h (**Fig. 2C-D**). These data support that NPY1R and NPY5R mRNA abundance is not only increased by hypoxia in general, but specifically via HIF-α stabilization.

**Figure 2.**
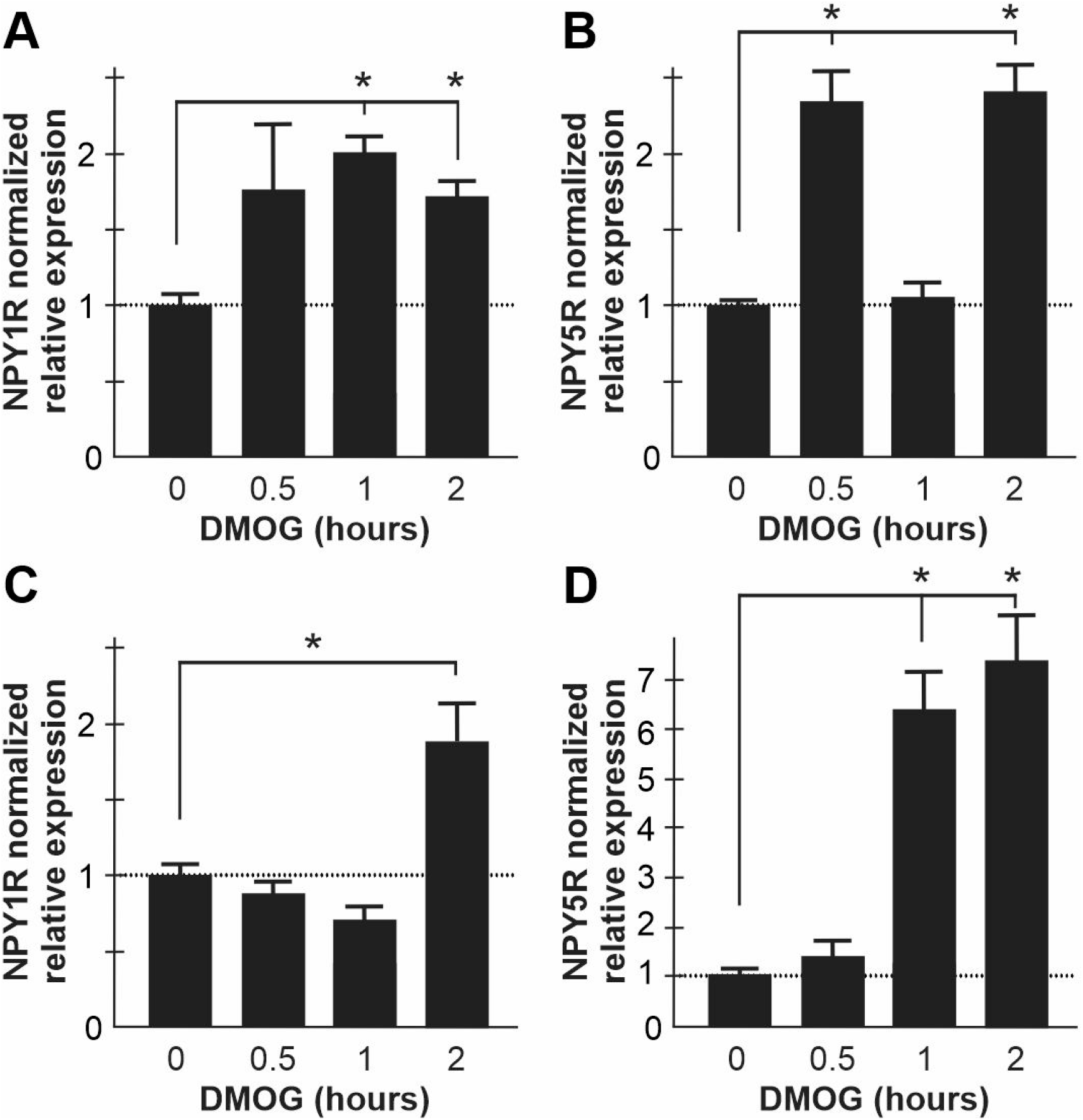
A hypoxia mimetic that stabilizes the HIFs increases the mRNA abundance of NPY1R and NPY5R. NPY1R and NPY5R mRNA abundance was measured in MCF7 (A-B) and MDA-MB-231 (C-D) breast cancer cells treated with a hypoxia mimetic, Dimethyloxalylglycine (DMOG), that is a cell permeable prolyl-4-hydroxylase inhibitor. Cells were lysed after 0.5, 1, and 2 h of DMOG treatment and mRNA abundance was measured by qRT-PCR. Data was made relative (dotted line) to DMSO vehicle control (time 0 h) and normalized to endogenous control genes *RPLP0* and *RPL13A*. Data (n = 3), mean ± s.e.m. * represents p < 0.05 using one-way ANOVA and Tukey’s HSD post-hoc test.

### Overexpression and inhibition of the HIFs affects NPY1R and NPY5R mRNA abundance

Exogenous HIF-α subunits were transfected into normoxic and hypoxic MCF7 and MDA-MB-231 cells to investigate whether they could drive NPY1R and NPY5R expression. These exogenous HIF-α subunits contain proline to alanine mutations that stabilize them in normoxia by escaping proteasomal targeting via prolyl hydroxylases (**Fig. S1**). In MCF7, NPY1R mRNA abundance significantly increased by 2.0 ± 0.07-fold in normoxia and by 2.27 ± 0.08-fold in hypoxia when exogenous stable HIF-2α was introduced relative to control (**Fig. 3A**). Exogenous stable HIF-1α significantly increased NPY1R mRNA abundance in MCF7 only in hypoxia by 1.48 ± 0.04-fold relative to control (**Fig. 3A**). NPY5R mRNA abundance significantly increased in normoxia by 1.67 ± 0.06-fold when stable HIF-1α was introduced, and in hypoxia by 2.94 ± 0.09-fold when stable HIF-2α was introduced relative to control (**Fig. 3B**). In normoxic MDA-MB-231 cells, introduction of both exogenous HIF-α subunits produced significant increases in both NPY1R (2.0 ± 0.06-fold for HIF-1α and 1.6 ± 0.04-fold for HIF-2α) and NPY5R (4.97 ± 0.31-fold for HIF-1α and 5.32 ± 0.08-fold for HIF-2α) mRNA abundance relative to control (**Fig. 3C-D**). In hypoxia, MDA-MB-231 cells displayed a significant increase only in NPY5R mRNA abundance by 3.51 ± 0.26-fold when exogenous stable HIF-1α was introduced relative to control (**Fig. 3D**). These data show that exogenous introduction of stable HIF-α subunits mostly increased the mRNA abundance of NPY1R and NPY5R.

**Figure 3.**
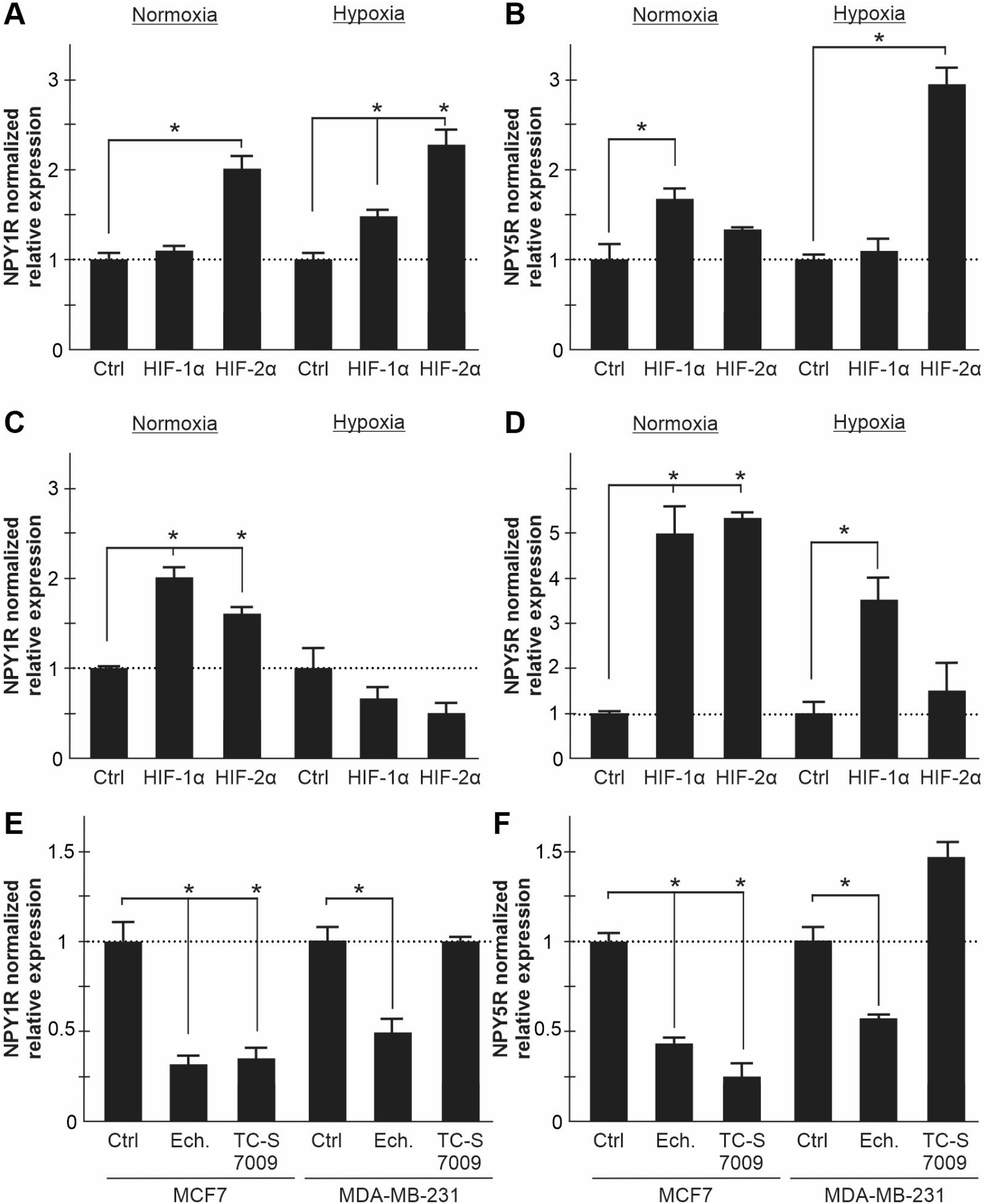
Overexpression and inhibition of the HIFs affects NPY1R and NPY5R mRNA abundance. (A-B) MCF7 and MDA-MB-231 (C-D) breast cancer cells were transfected with exogenous HIF-1α, HIF-2α, or their respective empty vector backbone control (Ctrl), and NPY1R (A and C) and NPY5R (B and D) mRNA levels were measured via qRT-PCR. Exogenous HA-HIF-α or FLAG-GFP-HIF-2α contain proline to alanine mutations that stabilize them in normoxia by escaping proteasomal targeting via prolyl hydroxylases. Data was made relative to the respective empty vector control (dotted line). (E) NPY1R and NPY5R (F) mRNA levels were measured via qRT-PCR in MCF7 and MDA-MB-231 cells treated with small molecule inhibitors specific for HIF-1 (Ech; Echinomycin) or HIF-2 (TC-S 7009) DNA binding activity. Data was made relative to an empty vehicle DMSO control (dotted line). All data (n=3) normalized to endogenous control genes *RPLP0* and *RPL13A*, and represented as mean ± s.e.m. * represents p < 0.05 using one-way ANOVA and Tukey’s HSD post-hoc test.

We next turned to pharmacological inhibition of endogenous HIFs. We treated hypoxic MCF7 and MDA-MB-231 cells with Echinomycin and TC-S 7009, selective inhibitors of HIF-1 and HIF-2, respectively. Cells were incubated in hypoxia for 24 h with or without Echinomycin or TC-S 7009. In hypoxic MCF7 cells, inhibition of HIF-1 reduced NPY1R and NPY5R mRNA levels to 0.32 ± 0.03 and 0.43 ± 0.02 the levels of the control, respectively (**Fig. 3E-F**). Similarly, inhibition of HIF-2 reduced hypoxic NPY1R and NPY5R mRNA levels to 0.35 ± 0.03 and 0.25 ± 0.04 the levels of the control, respectively (**Fig. 3E-F**). Treating hypoxic MDA-MB-231 cells with Echinomycin reduced NPY1R and NPY5R mRNA levels to 0.49 ± 0.04 and 0.57 ± 0.01 the levels of the control, respectively (**Fig. 3E-F**). Conversely, treatment of MDA-MB-231 cells with TC-S 7009 did not reduce the hypoxic mRNA levels of NPY1R or NPY5R (**Fig. 3E-F**). These data show that the HIFs are involved in the hypoxic induction of NPY1R and NPY5R.

### The HIFs bind to the promoters of *NPY1R* and *NPY5R* and upstream sequences

To test whether the HIFs are more directly implicated in the hypoxic induction of *NPY1R* and *NPY5R* transcription, we performed a Chromatin Immunoprecipitation (ChIP) assay. HIFs recognize Hypoxia Response Elements (HREs; 5’-RCGTG-3’) in the promoters of hundreds of genes to enhance their hypoxic expression (25). HREs are functionally activated when the downstream region contains an adjacent hypoxia ancillary sequence (HAS; 5’-CA(G|C)(A|G)(T|G|C)-3’) within 7 to 15 nucleotides (26). The *NPY1R* and *NPY5R* promoters and upstream regions were scanned for HREs and adjacent HASs. Several regions of interest fitting these criteria were identified, and primers were designed to amplify these genomic locations. Cells were transfected with stable exogenous HIF-1α, HIF-2α, or empty vector control followed by immunoprecipitation and RT-PCR. HIF-1α interacted with six of the seven genomic regions surveyed containing HREs and adjacent HASs within 15 kilobases of the *NPY1R* TSS in MDA-MB-231 (**Fig. 4A**) and four of the seven regions in MCF7 (**Fig. S2A**). Three sites containing HREs and adjacent HASs were identified within 15 kilobases upstream of the *NPY5R* TSS. All three sites were positive for HIF-1α interaction in MDA-MB-231 (**Fig. 4B**), and one of three sites were positive for HIF-1α association in MCF7 (**Fig. S2B**). The known HIF-1α target gene *BIRC5* (27) was used as a positive control (**Fig. 4C**). HIF-2α did not interact with any of the seven HRE/HAS-containing regions upstream of the *NPY1R* TSS in MDA-MB-231 (**Fig. 4D)**. However, HIF-2α did interact with the three genomic regions surveyed upstream of the *NPY5R* TSS (**Fig. 4E**) at comparable levels to the positive control gene *CITED2* (28) (**Fig. 4F**). Three of the seven sites upstream of the *NPY1R* TSS (**Fig. S2C**) and one of the three sites upstream of the *NPY5R* TSS (**Fig. S2D**) were positive for HIF-2α in MCF7. These data show that HIF-1α and HIF-2α directly interact (except HIF-2α upstream of the *NPY1R* TSS in MDA-MB-231) with the *NPY1R* and *NPY5R* promoter and upstream regions.

**Figure 4.**
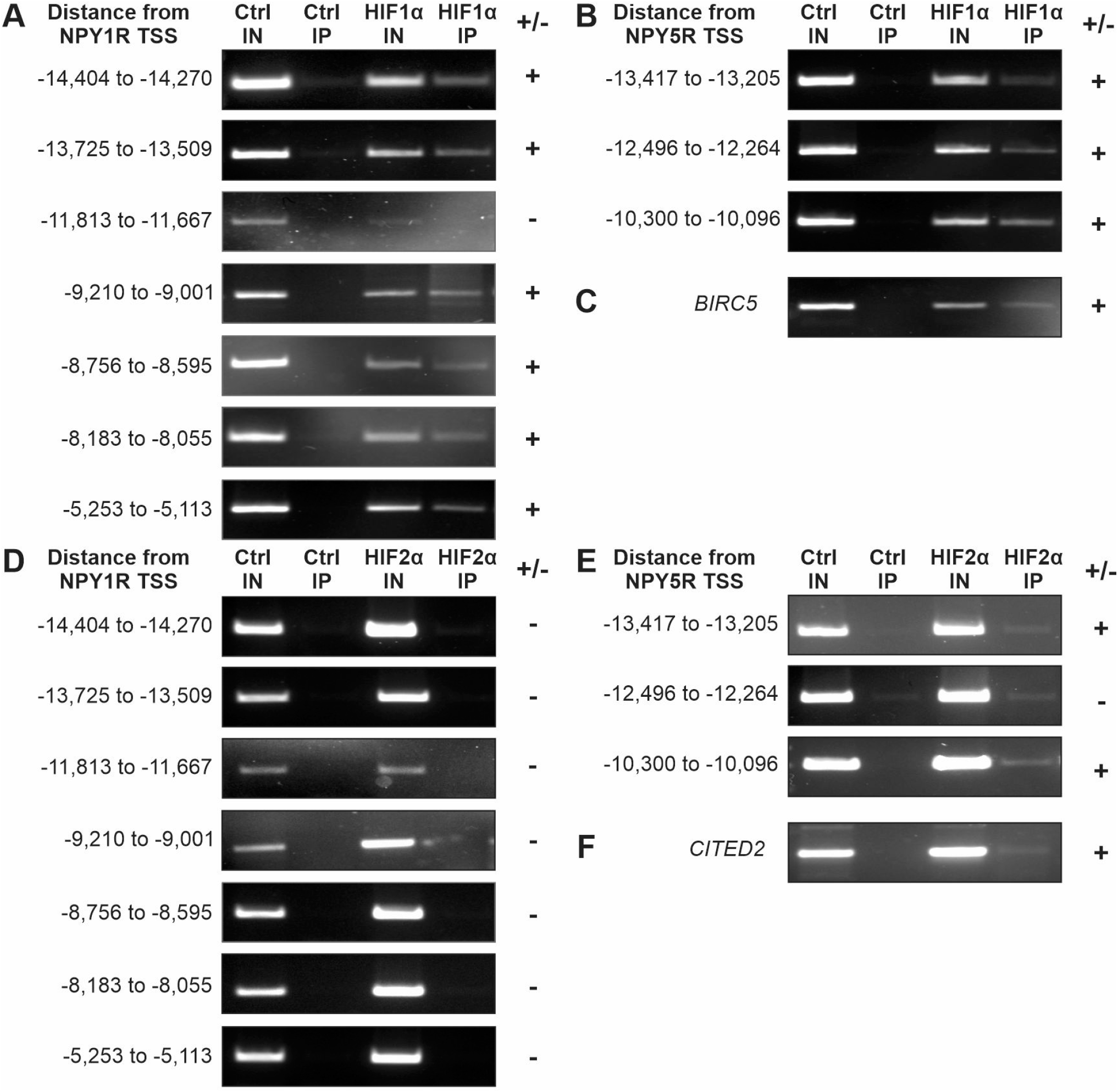
The HIFs bind to the promoters of *NPY1R* and *NPY5R* and upstream sequences. Chromatin Immunoprecipitation of exogenous HIF-1α (A-C) and HIF-2α (D-F) was performed in hypoxic MDA-MB-231 breast cancer cells. Genomic regions up to 15 kilobases upstream from the *NPY1R* and *NPY5R* transcription start sites (TSS) were scanned for Hypoxia Response Elements and downstream adjacent hypoxia ancillary sequences. Seven regions that fit these criteria were identified in the *NPY1R* TSS (A and D) and three regions in the *NPY5R* TSS (B and E). Primers were designed to amplify DNA regions via PCR that associated with either HIF-1α and HIF-2α. Control (Ctrl) antibody was a non-targeted IgG. Input (IN) was 10% whole cell lysate prior to immunoprecipitation. A genomic region was designated as positive (+) for HIF binding if the PCR amplified fragment was enriched in the HIF immunoprecipitation (IP) relative to the control. *BIRC5* (C) and *CITED2* (F) are two positive control genes that are known to have HIF-1α and HIF-2α binding sites, respectively, in their promoter regions.

### Stimulation of NPY1R and NPY5R induces MAPK signaling more rapidly in hypoxia

We next sought to determine whether the hypoxic induction of NPY1R and NPY5R had biological significance by measuring the activity of associated signaling cascades. NPY1R and NPY5R have been implicated in activating the MAPK pathway with subsequent increases in ERK1/2 phosphorylation (9,10). We stimulated normoxic and hypoxic MCF7 and MDA-MB-231 cells with either NPY (broad stimulation of NPYRs), NPY1R-specific ligand (Y1 agonist), or NPY5R-specific ligand (Y5 agonist). Hypoxic MCF7 cells responded more rapidly to NPY and Y5 agonist with a significant 1.75-fold and 5.25-fold higher pERK1/2, respectively, relative to normoxia at 5 min of stimulation (**Fig. 5A-B**). Conversely, stimulation of MCF7 cells with Y1 agonist produced a significant 2.57-fold higher pERK1/2 levels in normoxia relative to hypoxia at 5 min, but similar responses throughout the rest of the time course (**Fig. 5C**). Hypoxic MDA-MB-231 cells also responded more rapidly to NPY and Y5 agonist relative to normoxia with 3.28-fold and 1.94-fold higher pERK1/2 after 5 min of stimulation, respectively (**Fig. 5D-E**). Y1-specific stimulation of MDA-MB-231 cells produced little activation of ERK1/2 and similar responses in normoxia and hypoxia (**Fig. 5F**). These data suggest that hypoxic cells are primed to respond to NPY more rapidly than normoxic cells through NPY5R rather than NPY1R.

**Figure 5.**
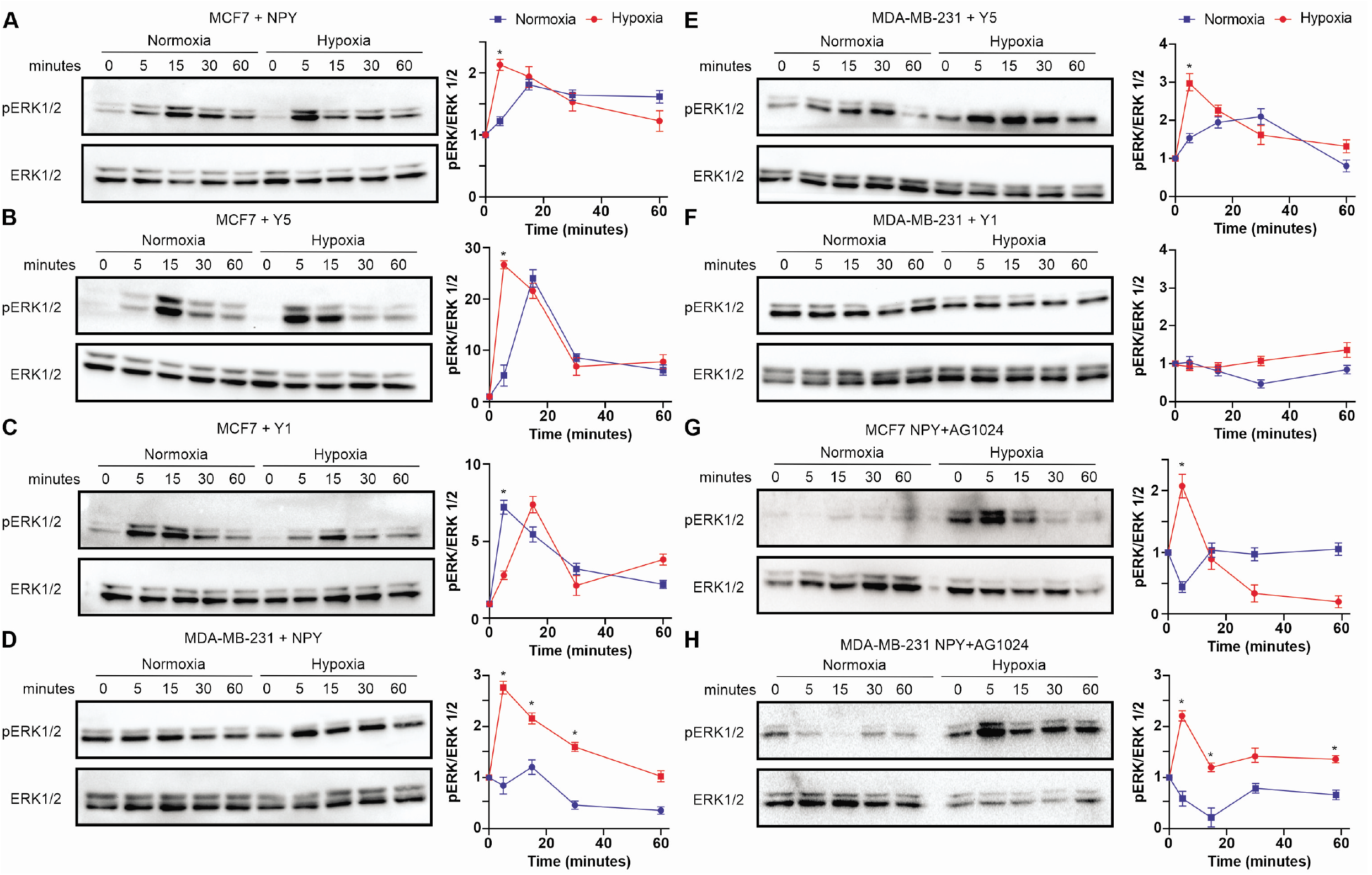
Stimulation of NPY1R and NPY5R induces MAPK signaling more rapidly in hypoxia. (A-C) MCF7 cells were treated with the general NPYR ligand NPY (A), NPY1R-specific Y1 agonist (B), and NPY5R-specific Y5 agonist (C) in normoxia (21% O_2_) and hypoxia (1% O_2_) for 5, 15, 30, and 60 min. MDA-MB-231 cells were treated with the NPY (D), Y1 agonist (E), and Y5 agonist (F) in normoxia and hypoxia for 5, 15, 30, and 60 min. (G-H) The IGF1R-specific inhibitor AG1024 was used to pre-treat MCF7 (G) and MDA-MB-231 (H) cells for 1 h before stimulating with NPY for 5, 15, 30, and 60 min. Activated ERK1/2 (pERK1/2) was normalized to total ERK1/2, quantified by densitometry, and made relative to time 0 for each condition. Data (n = 3), mean ± s.e.m. * represents p < 0.05 using one-way ANOVA and Tukey’s HSD post-hoc test.

Since NPYR can mediate the activation of ERK1/2 via transactivation with IGF1R in HEK293 cells (11), we tested whether IGF1R activity was required for NPY-dependent ERK1/2 phosphorylation in breast cancer cells. We pre-treated MCF7 and MDA-MB-231 cells with the specific IGF1R inhibitor AG1024 for 1 h, then stimulated with NPY to monitor ERK1/2 activation. In normoxic MCF7 cells, pERK1/2 peaked at 1.8 ± 0.17-fold induction at 15 min of NPY stimulation alone relative to time 0 (**blue line; Fig. 5A**). When AG1024 was present, normoxic NPY-dependent pERK1/2 induction was abolished with an initial decrease after 5 min that was 0.44 ± 0.08-fold relative to time 0 (**blue line; Fig. 5G**). In hypoxic MCF7 cells, AG1024 did not repress pERK1/2 induction, as similar peaks of 2.13 ± 0.09-fold and 2.07 ± 0.19-fold relative to time 0 were observed for NPY alone (**red line; Fig. 5A**) or with AG1024 (**red line; Fig. 5G**), respectively. In normoxic MDA-MB-231 cells, pERK1/2 displayed a mild induction that peaked at 1.2 ± 0.14-fold induction at 15 min of NPY stimulation alone relative to time 0 (**blue line; Fig. 5D**). In the presence of AG1024, normoxic NPY-dependent pERK1/2 induction was reversed with 0.2 ± 0.19-fold pERK1/2 levels after 15 min relative to time 0 (**blue line; Fig. 5H**). In the presence of AG1024, hypoxic MDA-MB-231 cells were still able to induce pERK1/2 with a peak of 2.21 ± 0.1-fold after 5 min of NPY stimulation relative to time 0 (**red line; Fig. 5H**), similar to the 2.76 ± 0.13-fold 5 min induction observed with NPY alone (**red line; Fig. 5D**). These data suggest that NPY-dependent stimulation of the MAPK/ERK pathway requires IGF1R activity in normoxia, but not in hypoxia.

### Hypoxic cells proliferate and migrate more than normoxic cells upon NPY stimulation

The phosphorylation of ERK1/2 stimulates cell proliferation and migration within the hypoxic tumor microenvironment (29,30). Therefore, we tested whether our observation of hypoxic MCF7 and MDA-MB-231 cells being more sensitive to NPY and Y5 stimulation produced more cell proliferation and migration in this context. NPY stimulation in MCF7 cells significantly increased cell migration in a scratch wound assay in both normoxia and hypoxia relative to their controls, but the increase was larger in hypoxia (2.29-fold) relative to normoxia (1.54-fold) (**Fig. 6A**). Further, only hypoxic MCF7 cells displayed an increase in cell migration when stimulated with Y5 agonist with a 1.88-fold increase relative to control (**Fig. 6A**). In MDA-MB-231, the only agonist that produced a significant increase in migration relative to the control was Y5 in hypoxia with a 1.63-fold change (**Fig. 6A**). Transwell assays produced similar findings to the scratch wound assay. In MCF7, NPY stimulation produced a significantly larger increase in migration in hypoxia (2.19-fold) than in normoxia (1.82-fold) relative to their controls (**Fig. 6B**). The Y5 agonist only produced an increase in migration in hypoxic MCF7 cells with a significant 2.49-fold change relative to control (**Fig. 6B**). None of the agonists produced an increase in MDA-MB-231 cell migration in the transwell assay beyond the effect of hypoxia alone (**Fig. 6B**). In both cell lines and both cell migration assays, the Y1 agonist did not produce an increase in migration but rather abolished the increase produced by hypoxia (**Fig. 6A-B**). Hypoxia significantly increased proliferation in both cell lines, but the agonists only produced further increases in hypoxia. NPY and Y5 stimulation increased proliferation by 1.5-fold and 2.35-fold, respectively, relative to hypoxic control **(Fig. 6C**). In MDA-MB-231 cells, Y1 and Y5 agonists produced proliferation increases of 1.6-fold and 1.63-fold, respectively, relative to hypoxic control (**Fig. 6C**). These data show that stimulation with NPY, likely through NPY5R, has a greater effect on cell migration in hypoxia. Moreover, stimulation with NPY produces more proliferation in hypoxic cells through both NPY1R and NPY5R.

**Figure 6.**
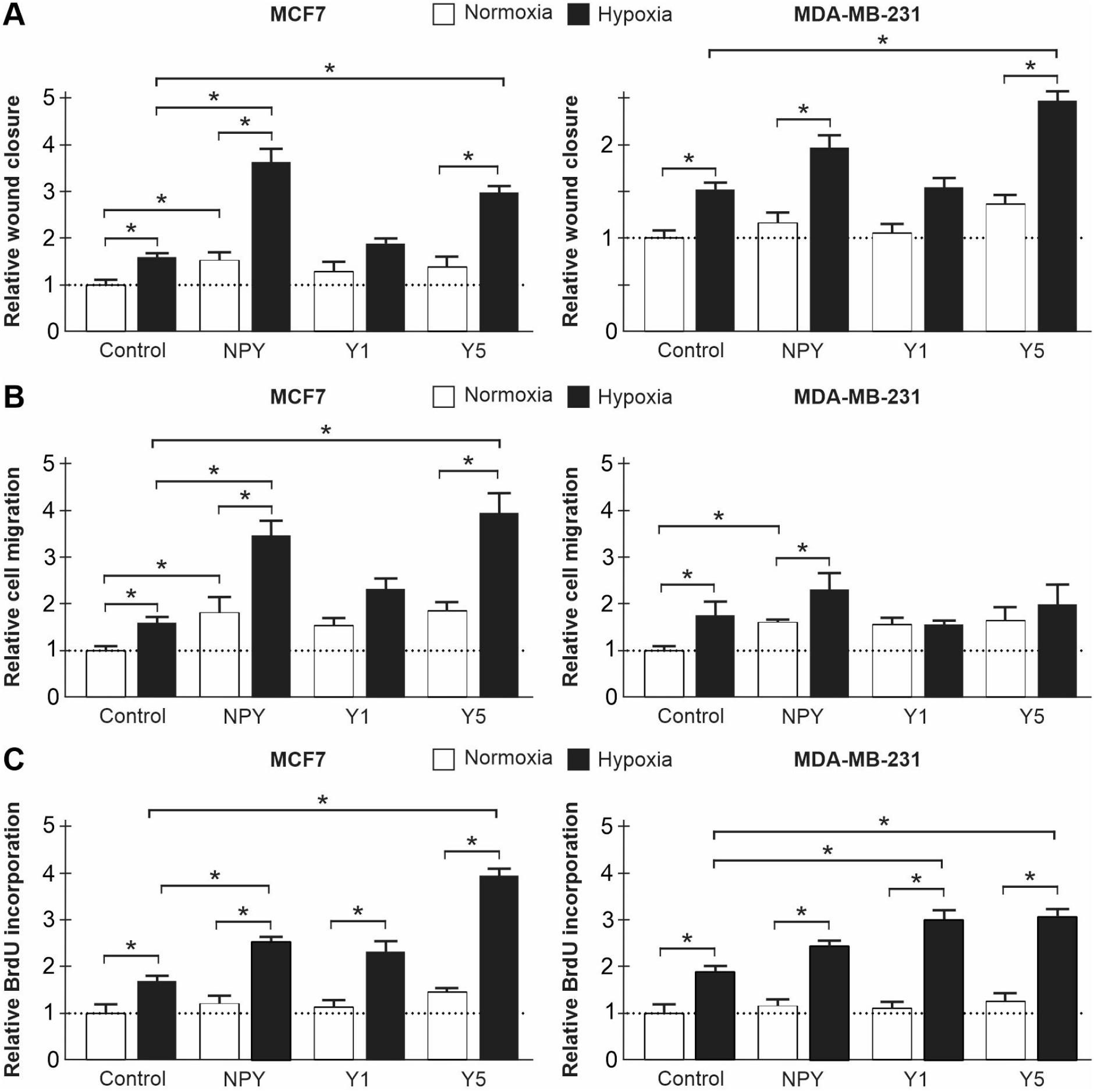
Hypoxic cells proliferate and migrate more than normoxic cells upon NPY stimulation. (A) A scratch wound was generated on a confluent monolayer of MCF7 and MDA-MB-231 cells that were stimulated with the general NPYR ligand NPY, NPY1R-specific ligand Y1 agonist, NPY5R-specific ligand Y5 agonist or vehicle control in normoxia (21% O2) and hypoxia (1% O2). Data (n=3) represents the mean % wound closure after 18 h (MCF7) or 6 h (MDA-MB-231) of stimulation by NPYR ligand relative to normoxic vehicle control. (B) Transwell migration assay in MCF7 and MDA-MB-231 cells stimulated with NPY, Y1 agonist, Y5 agonist or vehicle control in normoxia and hypoxia. Data (n=3) represents the number of cells that migrated after 18 h (MCF7) or 6 h (MDA-MB-231) of stimulation by NPYR ligand relative to normoxic vehicle control. (C) Cell proliferation measured in MCF7 and MDA-MB-231 cells stimulated with NPY, Y1 agonist, Y5 agonist or vehicle control in normoxia and hypoxia. Data (n=3) represents the mean % BrdU-positive cells after stimulation with NPYR ligand relative to normoxic vehicle control. Error bars, s.e.m. * represents p < 0.05 using one-way ANOVA and Tukey’s HSD post-hoc test.

## DISCUSSION

The NPYRs have received attention in the past couple decades because of their overexpression in a variety of human cancers. Breast cancer in particular has intrigued researchers because of its high frequency of NPYR overexpression and density compared to all other NPYR-positive tumors (16). This characteristic has been exploited to develop chemically modified analogs of NPY that are used in breast cancer imaging and diagnosis (31–33). Since breast tissue is highly innervated by the sympathetic nervous system, high local NPY concentration leads to heightened NPYR activity (9,10). Therefore, cancer therapeutics are also being aimed at the NPYRs.

We first examined whether NPYR expression in breast cancer cells was sensitive to changes in oxygen. Hypoxia is a feature of the tumor microenvironment that is also exploited by non-invasive imaging techniques to better treat, manage, and diagnose patients (34). Furthermore, hypoxia shifts NPY activity away from promoting cell death to drive cell proliferation, migration, and angiogenesis in Ewing sarcoma cells (24). We have directly implicated the HIFs in the hypoxic induction of NPY1R and NPY5R. The breast cancer cell lines in this study represent models of two different breast cancer subtypes: MCF7 is an Estrogen Receptor-positive model of the luminal A subtype and MDA-MB-231 is a triple negative breast cancer (TNBC) basal-like subtype (35). We performed a variety of assays to test whether both cell lines had similar hypoxic NPY1R and NPY5R, and whether both HIF-1α and HIF-2α were involved. The hypoxic induction of NPY1R and NPY5R displayed a temporal pattern in both cell lines (**Fig. 1A-D**) with increases after 1 h followed by a return to baseline levels. The exception was NPY1R in MDA-MB-231 cells that took 12 h to display a significant increase in mRNA abundance **(Fig. 1B**). The specific HIF-1α and HIF-2α inhibitors (Echinomycin and TC-S 7009, respectively) were potent at reducing NPY1R and NPY5R hypoxic induction except in MDA-MD-231 cells, which were insensitive to TC-S 7009 (**Fig. 3E-F**). Further, overexpressing stable HIF-2α had no effect on both NPY1R and NPY5R hypoxic induction in MDA-MB-231 cells, but did have an effect in normoxia where endogenous HIF levels are low (**Fig. 3C-D**). Overexpression studies are less elegant than inhibiting the endogenous protein, especially since endogenous HIFs saturate many of their binding sites in hypoxia (36). The basal breast cancer subtype, which includes MDA-MB-231, has an elevated baseline hypoxic gene signature relative to other subtypes (37). Moreover, TNBC has a high baseline HIF pathway activation, and high endogenous HIF levels in normoxia possibly due to a paracrine signaling mechanism that involves glutamate secretion (38,39). Indeed, we observed a lower hypoxic induction of HIF-2α in MDA-MB-231 compared to MCF7 cells, possibly due to higher baseline HIF-2α levels in normoxic MDA-MB-231 cells (**Fig. S1**). Our data support that the HIFs are involved in the hypoxic induction of NPY1R and NPY5R, and that HIF-1α is the dominant homolog in basal subtypes of TNBC.

A HIF-dependent induction of NPY1R and NPY5R does not indicate that the HIFs are directly involved. For example, a HIF-dependent gene could initiate a chain of events leading to increased NPY1R and NPY5R expression. The HIFs bind to HREs, but these core sequences of 5’-RCGTG-3’ vastly outnumber the 50-100 validated HIF target genes to permit prediction of HIF binding (25). HIF binding has often been determined through conservation in mammals and proximity to the TSS, but this over-represents short distance interactions (40). The identification of HIF enhancer regions 7 to 15 nucleotides downstream of HREs, termed HAS, have narrowed strong candidate HIF binding sites (26). An unbiased analysis of HIF binding across the whole genome through ChIP and next-generation high-throughput sequencing showed that HIFs can bind to DNA far from the TSS. Over 60% of HIF-1α and 80% of HIF-2α binding sites were more than 2.5 kb from the TSS, with some even greater than 100 kb (41). The range of HIF binding sites identified through ChIP in our study were between 5-15 kb from the *NPY1R* and *NPY5R* TSS (**Fig. 4**). MDA-MB-231 displayed more HIF binding in these regions than MCF7, and a preference for HIF-1, which is consistent with basal-type TNBC having a more active HIF pathway (37–39) and a preference for HIF-1α (**Fig. 3C-F and Fig. S1**). There is evidence in the literature that the HIFs can function at long genomic intervals, but we cannot rule out that NPY1R and NPY5R expression is indirectly induced by HIF-regulated networks.

Hypoxia sensitized cells to NPY stimulation where pERK1/2 levels peaked earlier than in normoxic cells. This is especially evident in MDA-MB-231, an aggressive cell line with high baseline pERK (42), where normoxic NPY stimulation produced modest changes in pERK1/2 levels compared to significant induction in hypoxia (**Fig. 5D-F**). Furthermore, of the isotype-specific agonists, only the Y5 agonist demonstrated hypoxic sensitization (**Fig. 5B and E**) suggesting that NPY5R has a greater role in hypoxia. To further explore the mechanism, we show that IGF1R activity is involved in NPY-mediated activation of MAPK/ERK in normoxic cells, but not in hypoxic cells (**Fig. 5G-H**). Hypoxic cells were also more responsive to NPY and Y5 agonist than normoxic cells with respect to cell migration and proliferation, both outcomes of active MAPK/ERK (**Fig. 6**). The Y1 agonist negated the hypoxic increase in cell migration in both cell lines (**Fig. 6A-B**) and did not increase cell proliferation in MCF7 compared to its hypoxic control (**Fig. 6C**). Indeed, NPY1R stimulation can be inhibitory, including toward cell growth in MCF7 (43). Our data suggest that hypoxic cells are more sensitive than normoxic cells to NPY mostly through NPY5R. This could be due to the increased levels of NPY1R and NPY5R in hypoxia. IGF1R is also a hypoxia-induced protein (44), which could affect the stoichiometry with its inhibitor AG1024, reducing the effect of this chemotherapeutic on hypoxic cells. Since hypoxia affects drug metabolism and pharmacokinetic capacity (45), this should also be considered as a possible mechanism. In breast cancer, NPY stimulation of NPY5R could have a prominent role in hypoxia-driven metastasis.

A growing body of evidence suggests that neuroendocrine factors contribute to the initiation, development, and progression of breast cancer. NPY contributes to proliferation (9,10) migration (9,10) and angiogenesis (18), however NPYR activity in hypoxia had yet to be examined in a breast cancer model. Our findings that NPYRs are sensitive to changes in oxygen support the postulate that NPY in a low oxygen microenvironment may contribute to tumor progression and metastasis. Questions remain whether tumor neuronal release of NPY or NPY proteolytic activity are affected by hypoxia as reported in other models (24). Future *in vivo* studies may help elucidate the effects of hypoxia on ligand, proteolytic activity, receptors, and the functional impact of the NPY system within the tumor microenvironment. Our data highlight NPY1R and NPY5R as hypoxia-inducible and HIF-dependent, rendering hypoxic cells more sensitive to NPY stimulation. This study should inform the development and use of NPYR antagonists in breast cancer therapy whereby targeting the NPY5R could exploit a vulnerability in hypoxic cells, which are more metastatic (20).

## EXPERIMENTAL PROCEDURES

### Cell Culture, Cell Lines, and Pharmacological treatments

MDA-MB-231 and MCF7 cells were obtained from the from the American Type Culture Collection and maintained as suggested. These were tested for mycoplasma contamination and characterized by short tandem repeat and Q-band assays. Normoxic cells were maintained in a humidified chamber (ambient O_2_, 5% CO_2_, and 37 °C). Hypoxic cells were incubated in a HypOxystation H35 (HypOxygen, Frederick, MD, U.S.A.) at 1% O_2_, 5% CO_2_, and 37 °C. The hypoxia mimetic DMOG was used at 1 mM for 0.5 h, 1 h and 2 h in normoxia. The HIF-1α inhibitor Echinomycin (20 nM) and the HIF-2α inhibitor TC-S 7009 (100 μM) were given to cells before a 24 h hypoxic incubation. The IGF1R inhibitor AG1024 (MedChemexpress; 10 μM) was given to cells 1 h prior to treatment with NPYR agonists (Tocris; 10 nM): NPY (cat#1153), NPY1R-specific (Cat#1176), or NPY5R-specific (cat#1365).

### RNA Isolation and qRT-PCR

RNA was extracted using RiboZol (VWR) per the manufacturer’s instructions. RNA (2 μg) was reverse transcribed using a high-capacity cDNA reverse transcription kit (Applied Biosystems). Primers (5’ to 3’) were as follows: NPY1R, CCATCGGACTCTCATAGGTTGTC (forward) and GACCTGTACTTATTGTCTCTCATC (reverse); NPY5R, CCTCAGGTGAAACTCTCTGGCA (forward) and GAGAAGGTCTTTCTGGAGCAGG (reverse); RPLP0, AACATCTCCCCCTTCTCC (forward) and CCAGGAAGCGAGAATGC (reverse). Quantitative PCR performed using SsoAdvanced Universal SYBR Green Supermix (BioRad). Data was analyzed with CFX manager software (Bio-Rad). Relative fold change in expression was calculated using the ΔΔCT method, and transcript levels were normalized to endogenous controls *RPLP0* and *RPL13A*.

### Western Blotting

Standard western blotting procedure was used. Primary antibodies: NPY1R (Abcam, NPY1R11-A), NPY5R (Abcam, NPY5R11-A), HIF-1α (Novus Biological, NB100-123), HIF-2α (Novus Biological, NB100-122), Phospho-p44/42 MAPK (T202/Y204) (Cell Signaling, 4370S), p44/42 MAPK (Erk1/2) (Cell Signaling, 4695S), GAPDH (Cell Signalling, 5174S). HRP-conjugated antibodies: Promega anti-Rabbit IgG (W401B) and anti-mouse IgG (W402B). Densitometry was performed using Image Lab (BioRad).

### DNA constructs and Transfection

Transient HIF-1α and HIF-2α expression was performed using the following expression vectors: FLAG–GFP–HIF-2α in a pAdlox backbone was a gift from Stephen Lee (Miami, FL), and HA– HIF-1α was a gift from William Kaelin (Addgene plasmid 18955, RRID:Addgene_18955 [http://n2t.net/addgene:18955]. These vectors produce stable non-degradable proteins that can be expressed in normoxia and retain their function. Cells were transfected with 4 μg DNA complexed with 10 μl Lipofectamine 2000 (Invitrogen) diluted in 500 μl of serum free DMEM and added to cells at 37 °C for 8 h followed by replenishment with complete medium. Transfected cells were incubated in normoxia or hypoxia for 24 h before isolating protein and RNA.

### Chromatin immunoprecipitation (ChIP)

MCF7 and MDA-MB-231 cells were transfected with empty vector control, HA-HIF-1ɑ or FLAG-GFP-HIF-2ɑ. Transfection was followed by 24 h of hypoxia. Samples were subjected to protein-DNA cross-linking with 1% paraformaldehyde for 10 min. Cells were washed with 1XPBS and fixation was arrested using 125 mM glycine. 1% phenylmethylsulfonyl fluoride in 1XPBS was applied to cells and then pelleted. Hypotonic lysis buffer (10 mM Tris-HCl, pH 8.1, 10 mM KCl, 2 mM MgCl2, 2.5 mM sodium pyrophosphate, 1 mM beta-glycerophosphate, 1 x protease inhibitor cocktail (NEB), 2 mM *N*-ethylmaleimide (NEM), 2 mM sodium orthovanadate (NaO), 2 mM sodium fluoride (NaF), 1 mM dithiothreitol) was added on ice for 10 min to release nuclei. 10% IGEPAL was added to the solution and the sample was vortexed and incubated on ice briefly. The pellet was resuspended in nuclear lysis buffer (50 mM Tris-HCl, pH 8.1, 10 mM EDTA, 1% sodium dodecyl sulfate (SDS), 1 x protease inhibitor cocktail, 2 mM NEM, 2 mM NaO, 2 mM NaF). Samples were sonicated and cellular debris removed via centrifugation. Supernatant was diluted with ChIP dilution buffer (10 mM EDTA, 1% Triton X-100, 150 mM NaCl, 20 mM Tris-HCl, pH 8.1, 1 x protease inhibitor cocktail, 2 mM NEM, 2 mM NaO, 2 mM NaF). Magnetic beads (Cell Signaling), Anti-HA Magnetic Beads (Pierce), and Anti-FLAG M2 Magnetic Beads (Sigma) were blocked with 2% bovine serum albumin, suspended in TBS, and then lysate was applied to the beads. Following overnight incubation, beads were washed subsequently in low-salt immune complex wash buffer (0.1% SDS, 1% Triton X-100, 2mM EDTA, 20 mM Tris-HCl, pH 8.1, 150 mM NaCl), high-salt immune complex wash buffer (0.1% SDS, 1% Triton X-100, 2mM EDTA, 20 mM Tris-HCl, pH 8.1, 500 mM NaCl), lithium chloride immune complex wash buffer (0.25 mM LiCl, 1% IGEPAL NP-40, 1% sodium deoxycholate, 1 mM EDTA, 10 mM Tris-HCl, pH 8.1), and a Tris-EDTA wash buffer (10 mM Tris-HCl, pH 8.1 and 1 mM EDTA). The bound complexes of interest were then eluted using SDS buffer (1% SDS and 50 mM NaHCO_3_). Crosslinking was reversed with 2.5M NaCl at 65 °C for 2.5 h. Samples were then treated with 1 mg/mL of RNase at 65 °C for 2.5 h, followed by 0.5M EDTA, 1M Tris-HCl, pH 6.5 and 10 mg/mL Proteinase K at 37 °C for 1 h. Finally, samples were purified using the GenepHlow Gel/PCR Kit (Geneaid) and separated on an agarose gel via electrophoresis. Histone H3 (Cell Signaling) antibody (2 μg/ml) was used as a positive control. Primers were designed to amplify short *NPY1R* and *NPY5R* promoter region amplicons using PCR (**Table S1**). Known HIF-1 target CITED2 (forward - CAAGTCAATGAACCAAACGG and reverse - ATAGATAACGTGGTAATCGC) and HIF-2 target survivin (forward 5’-GCGTTCTTTGAAAGCAGT-3’ and reverse 5’-ATCTGGCGGTTAATGGCG-3’) were used as positive controls. A positive

### Cell migration assays

For wound closure assays, 10^6^ cells were seeded in a 6 cm plate and grown to 80% confluence. Cells were washed with PBS and incubated with serum-reduced (0.5% FBS) media overnight at 37 °C in normoxia or hypoxia. Five scratches were made using a P200 pipette tip, followed by a PBS wash. Cells were incubated with serum-reduced media supplemented with Insulin-Transferrin-Selenium (Gibco). Treatment media containing 10 nM agonists NPY, NPY1R-specific, or NPY5R-specific was then added to plates that were incubated in either normoxia or hypoxia. Wounds were imaged at 0 h and 18 h for MCF7 or 6 h for MDA-MB-231 post treatment using the Ti-S microscope (Nikon) and analyzed with NIS-Elements software (Nikon).

For transwell migration assays, 12 well inserts with 8 μm pores (BD Biosciences) were used to assess cell chemotaxis. After 24 h of serum starvation, 5×10^4^ cells were seeded in the upper chamber in serum-free media. Peptidergic (10 nM agonist)-treated media (NPY, NPY1R-specific, or NPY5R-specific) was added to the bottom chamber. After 18 h (MCF7) or 6 h (MDA-MB-231) of incubation in normoxia and hypoxia, non-migrated cells were scraped from the membrane with a cotton swab, and migrated cells were fixed in methanol and stained with Hoescht. The membranes were removed, mounted on slides, and imaged on a TiS microscope (Nikon). Migrated cells were quantified using Matlab© software by a blinded experimenter as previously described (46).

### BrdU-ELISA

Cells (5×10^3^) were seeded in a 96-well plate followed by 24 h of serum starvation. Media was replaced with serum reduced (0.5% FBS) treatment media containing NPY, NPY1R-specific, or NPY5R-specific agonists at 10 nM. Cells were incubated at 37 °C in normoxia or hypoxia for 20 h and then with 1X BrdU substrate for an additional 4 h. The substrate is from the BrdU Cell Proliferation ELISA Kit (Abcam) and the manufacturer’s instructions were subsequently followed. The plates were read using a spectrophotometric microtiter plate reader at 450 nm.

### Statistical Analyses

Statistical analyses were performed using GraphPad and data are presented as mean ± s.e.m. Statistical differences between treatments were evaluated by one-way ANOVA followed by Tukey’s HSD test.

## SUPPORTING INFORMATION

This article contains supporting information.

## ACKNOWLEDGMENTS

We thank Erin Specker and Colin Jamieson for their technical assistance. This work was funded by the Ontario Ministry of Research and Innovation and the Natural Sciences and Engineering Council of Canada grant number 04807 to J.U.

## AUTHOR CONTRIBUTIONS

PJM, SP, and SK conducted all the experiments and analyzed most of the results. BKA analyzed and quantified the cell migration data. PJM, SP, SK, and JU wrote the paper. JU supervised and funded the project.

## CONFLICT OF INTEREST

The authors declare that they have no conflicts of interest with the contents of this article.

## Notes

### Competing Interest Statement

The authors have declared no competing interest.

