## Supplemental Information for "HIF-dependent Neuropeptide Y Receptor Y1 and Y5 expression sensitizes hypoxic cells to NPY stimulation"

Supporting Information:

Table S1

Figure S1

Figure S2

| Gene name | Distance from Transcription Start Site (TSS) | Forward (5' to 3') | Reverse (5' to 3') |
|---|---|---|---|
| *NPY 1R* | -14,404 to -14,270 | GGAGTTTCATGGGGCATTTAAGAAGG | GGATAAGGCCCTTAGCCCAAAGCC |
| *NPY 1R* | -13,725 to -13,509 | CCTTCCTCAAACTTAGTGGAATACAACTCATGG | CTAACTTTCAAGTCCCTCCTTTTGGCAATAG |
| *NPY 1R* | -11,813 to -11,667 | GGCGTCAAAGATCCAAAGAAATGCAG | GACATTGATAAGGGGGTAATTCCATG |
| *NPY 1R* | -9,210 to -9,001 | CAGTGGGCCAAAGAAAGGAGG | CAGCCCAAACCAGGAATCAAAG |
| *NPY 1R* | -8,756 to -8,595 | CACCTTTGTGAGGTGTTTGTGG | GAAGATCCGCTGGTATTTCTCCTGCTC |
| *NPY 1R* | -8,183 to -8,055 | CGAGTTCGCCTATCCCACACCC | CAAGCCCACCCTTCTCCGGCTCAT |
| *NPY 1R* | -5,253 to -5,113 | CATCGGAGTTCTAATCAGGGAACTC | CAGTCTACCTGGGAACCAATGGC |
| *NPY 5R* | -13,417 to -13,205 | CTCCCTCAAGCCAGAACCTAAGAATTGTC | CTGGGAAGCTGTATGACTGGATG |
| *NPY 5R* | -12,496 to -12,264 | CTTATGTGTTAGCTCTGCCTCCTC | CTGTTCAAAGCATAAGCTCAGGACC |
| *NPY 5R* | -10,300 to -10,096 | TAACTTGTCCACTCCAAGTGTTC | GAACCTTTAGGTTGAAGTCACTG |

Table S1. Primers used to detect genomic regions upstream of the *NPY1R* and *NPY5R* transcription start sites (TSS) after Chromatin Immunoprecipitation of HIF-1α and HIF-2α (Fig. 4 and Fig. S2).

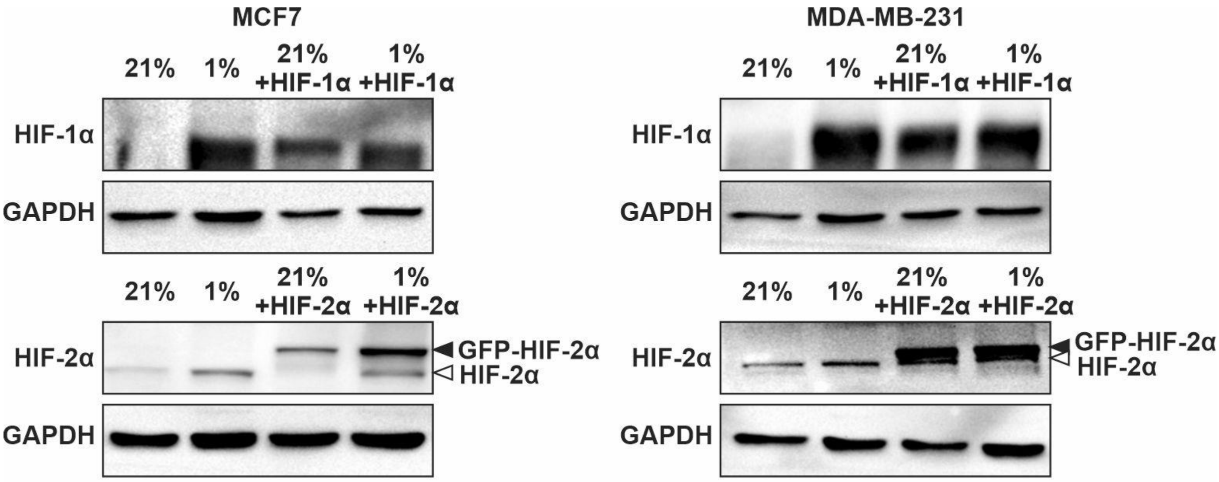


**Figure S1. Endogenous and exogenous HIF-1α and HIF-2α in transfected MCF7 and MDA-MB-231 cells.** Cells were transiently transfected with HA-HIF-1α and FLAG-GFP-HIF-2α and incubated in normoxia (21% $O_2$) or hypoxia (1% $O_2$). GAPDH used as a loading control.

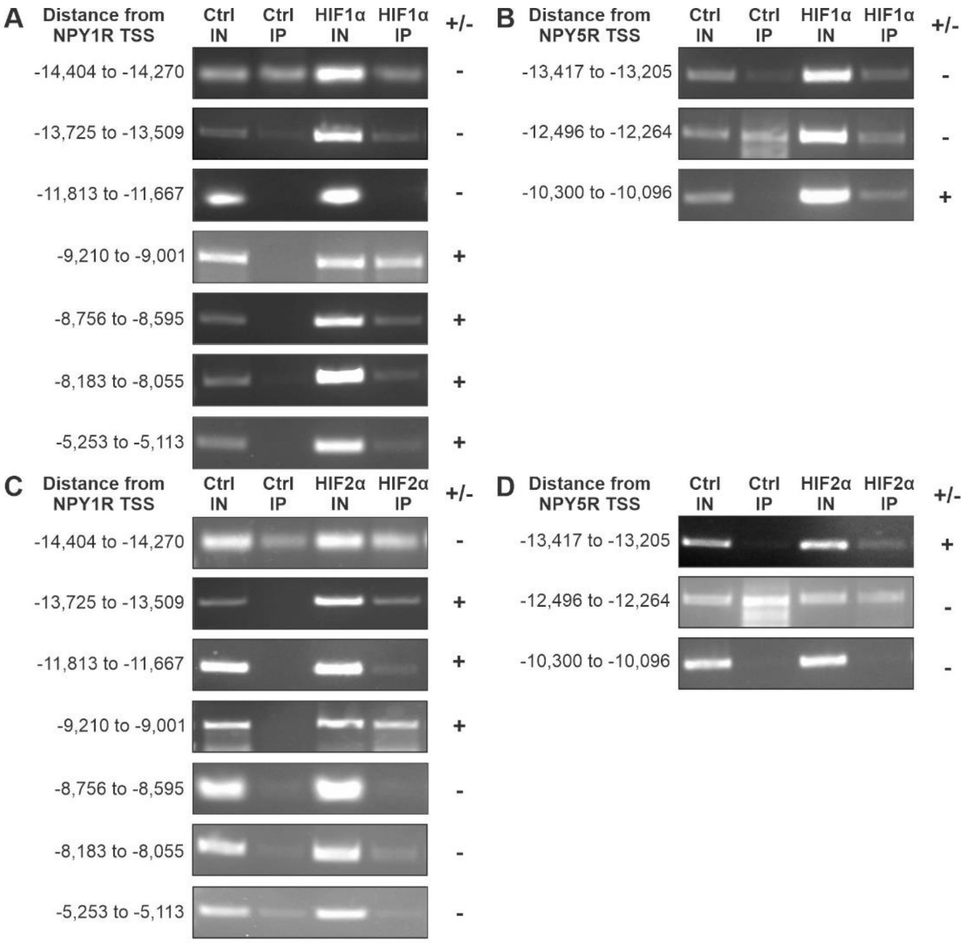


**Figure S2. The HIFs bind to the promoters of *NPY1R* and *NPY5R* and upstream sequences.** Chromatin Immunoprecipitation of exogenous HIF-1α (A-B) and HIF-2α (C-D) was performed in hypoxic MCF7 breast cancer cells. Genomic regions up to 15 kilobases upstream from the *NPY1R* and *NPY5R* transcription start sites (TSS) were scanned for Hypoxia Response Elements and downstream adjacent hypoxia ancillary sequences. Seven regions that fit these criteria were identified in the *NPY1R* TSS (A and C) and three regions in the *NPY5R* TSS (B and D). Primers were designed to amplify DNA regions via PCR that associated with either HIF-1α and HIF-2α. Control (Ctrl) antibody was a non-targeted IgG. Input (IN) was 10% whole cell lysate prior to immunoprecipitation. A genomic region was designated as positive (+) for HIF binding if the PCR amplified fragment was enriched in the HIF immunoprecipitation (IP) relative to the control.
